# Phylogenomic analysis clarifies the evolutionary origin of *Coffea arabica* L

**DOI:** 10.1101/2020.03.22.002337

**Authors:** Yves Bawin, Tom Ruttink, Ariane Staelens, Annelies Haegeman, Piet Stoffelen, Jean-Claude Ithe Mwanga Mwanga, Isabel Roldán-Ruiz, Olivier Honnay, Steven B. Janssens

**Affiliations:** Flanders Research Institute for Agriculture, Fisheries, and Food (ILVO), Belgium; Plant Conservation and Population Biology, KU Leuven, Belgium; Crop wild relatives and useful plants, Meise Botanic Garden, Belgium; Department of Plant Biotechnology and Bioinformatics, Ghent University, Belgium; Centre de Recherche en Sciences Naturelles (CRSN), D.R. Congo

**Keywords:** Allopolyploidy, *Coffea arabica* (Arabica coffee), genotyping-by-sequencing, hybridization, molecular dating, self-compatibility

## Abstract

Interspecific hybridization events have played a major role in plant speciation, yet, the evolutionary origin of hybrid species often remains enigmatic. Here, we inferred the evolutionary origin of the allotetraploid species *Coffea arabica*, which is widely cultivated for Arabica coffee production.

We estimated genetic distances between *C. arabica* and all species that are known to be closely related to *C. arabica* using genotyping-by-sequencing (GBS) data. In addition, we reconstructed a time-calibrated multilabeled phylogenetic tree of 24 species to infer the age of the *C. arabica* hybridization event. Ancestral states of self-compatibility were also reconstructed to infer the evolution of self-compatibility in *Coffea*.

*C. canephora* and *C. eugenioides* were confirmed as the putative progenitor species of *C. arabica.* These species most likely hybridized between 1.08 million and 543 thousand years ago.

We inferred the phylogenetic relationships between *C. arabica* and its closest relatives and shed new light on the evolution of self-compatibility in *Coffea*. Furthermore, the age of the hybridization event coincides with periods of environmental upheaval, which may have induced range shifts of the progenitor species that facilitated the emergence of *C. arabica*.

## Introduction

Interspecific hybridization events have played a major role in plant speciation (Mallet, 2005; Whitney *et al*., 2010). Most known hybrid species, including wheat (*Triticum* spp.), cotton (*Gossypium* spp.), and cabbage (*Brassica* spp.), are allopolyploids, which are hybrids with an increased chromosomal content compared to their diploid progenitor species (Soltis & Soltis, 2009; Renny-Byfield & Wendel, 2014). Because many allopolyploids became ecologically divergent or geographically isolated from their closest relatives, inference of their ancestry solely based on non-molecular characteristics is often difficult (Abbott *et al*., 2013). The genome of allopolyploid species consists of different subgenomes, each originating from one of its progenitor species. Even though subgenomes may lose genomic segments via a process called ‘biased fractionation’, ancestral polymorphisms between progenitor species remain present throughout the extant allopolyploid genome and can provide crucial information about the progenitor species (Pelé *et al*., 2018; Wendel *et al*., 2018).

*Coffea arabica* L. is the only known natural allopolyploid species in the genus *Coffea* (2n = 4x = 44) and one of the few known self-compatible species within its genus (Charrier & Berthaud, 1985). Cultivated across the tropics and subtropics, *C. arabica* is one of the most valuable agricultural commodities, accounting for about 60% of the global coffee production (ICO, 2020a,b). Nowadays, wild *C. arabica* populations are only found in the Afromontane rainforests of southwest Ethiopia, although small isolated populations also occur in Northern Kenya and the Eastern part of South Sudan (Davis *et al*., 2006). Wild *C. arabica* populations are currently threatened by climate change (Davis *et al*., 2012; Moat *et al*., 2019), increasing pest and disease pressure (Hindorf & Omondi, 2011; Vega *et al*., 2015), deforestation (Tadesse *et al*., 2014; Geeraert *et al*., 2019), and introgression of cultivar alleles into wild individuals (Aerts *et al*., 2012, 2017). Wild coffee species, including wild *C. arabica*, carry valuable genetic resources for coffee breeding. However, more than half of the currently described 125 wild *Coffea* species are threatened with extinction (Davis *et al*., 2019; Govaerts *et al*., 2020). Because many of these species are also poorly conserved in *ex situ* collections, the decline of wild *Coffea* species is a widely acknowledged problem (Davis *et al*., 2019; Moat *et al*., 2019).

The species *C. arabica* emerged through the natural hybridization of two *Coffea* species followed by a whole genome duplication, probably during a single allopolyploidization event (Clarindo & Carvalho, 2008; Lashermes *et al*., 2014; Scalabrin *et al*., 2020). The current geographical range of *C. arabica* does not overlap with that of any other *Coffea* species so that geographical co-existence cannot be used to put forward candidate progenitors (Davis *et al*., 2006). Using genomic *in-situ* hybridization (GISH) and Restriction fragment length polymorphism (RFLP) markers, *C. canephora* Pierre ex A.Froehner and *C. eugenioides* S.Moore have been identified as the closest extant relatives of *C. arabica* (Lashermes *et al*., 1999). Although cytogenetic methods such as GISH are considered reliable for studying hybridization (Chester *et al*., 2010), a certain ambiguity remains regarding the progenitor species of *C. arabica*. Based on GISH and fluorescence in-situ hybridization (FISH), Raina *et al*. (1998) suggested *C. congensis* A.Froehner as progenitor species of *C. arabica* instead of *C. canephora*. Hamon *et al*. (2009), however, could not discriminate between *C. canephora* and *C. congensis* as putative progenitor of *C. arabica* using FISH and fluorochrome banding (CMA, DAPI). Moreover, genetic divergence in plastid DNA regions or in the internal transcribed spacer (ITS) sequence were too low to resolve phylogenetic relationships between *C. arabica* and other *Coffea* species (Berthou *et al*., 1980, 1983; Lashermes *et al*., 1997; Cros *et al*., 1998; Maurin *et al*., 2007; Tesfaye *et al*., 2007). In addition, species such as *C. anthonyi* Stoff. & F.Anthony, *C. heterocalyx* Stoff., and *C. kivuensis* Lebrun which are closely related to *C. eugenioides* and which share some key traits with *C. arabica*, have not been consistently included in evolutionary studies of *C. arabica.* The habitus of *C. kivuensis* is very similar to that of *C. arabica* and both species have similar leaf, flower, and fruit characteristics. Furthermore, *C. heterocalyx* and *C. anthonyi* are, together with *C. arabica*, the only known self-compatible species in *Coffea* (Coulibaly *et al*., 2002; Stoffelen *et al*., 2009). Taken together, and despite all these research efforts, the origin of *C. arabica* remains elusive. The unambiguous inference of the phylogenetic relationships between *C. arabica* and its relatives remains crucial to understand the evolutionary history of these species (Tesfaye *et al*., 2007; Stoffelen *et al*., 2009).

Estimating the age of the interspecific hybridization event at the origin of *C. arabica* has been the purpose of several studies. Based on the frequency of synonymous substitutions in a phosphoenolpyruvate carboxylase kinase gene of *C. canephora* and its orthologue in the corresponding *C. arabica* subgenome, Yu *et al*. (2011) estimated the divergence time between *C. canephora* and *C. arabica* around 665 000 years ago. Cenci *et al*. (2012) calculated the minimum age of *C. arabica* between 10 000 and 50 000 years by comparing the substitution frequency between the *C. arabica* subgenomes in a 50 kilobase region that was assumed to be duplicated in *C. arabica* after its emergence, with the substitution frequencies in other regions of the *C. arabica* genome. Furthermore, it has very recently been shown that genetic diversity levels in simulated populations of *C. arabica* became similar to those observed in *C. arabica* accessions when the origin of *C. arabica* was set to 10 000 years, again supporting a very recent origin of the species (Scalabrin *et al*., 2020). Given these divergent estimates, a more elaborate molecular dating analysis is still needed (Yu *et al*., 2011; Cenci *et al*., 2012).

The availability of affordable genome-wide sequencing techniques now enables the reconstruction of hybridization events based on subtle differences in genome sequence between hybrids and their relatives (Payseur & Rieseberg, 2016). For example, genotyping-by-sequencing (GBS) has been used to identify the origin of hybrid species with high accuracy in soybean (*Glycine* spp.) and vanilla (*Vanilla* spp.) (Sherman-Broyles *et al*., 2017; Hu *et al*., 2019). GBS markers may contain more information about the evolutionary history of a species than single gene sequences because they originate from a large number of regions located across the genome. The value of GBS to reconstruct phylogenetic relationships within the genus *Coffea* has been demonstrated by Hamon *et al*. (2017) and Guyeux *et al*. (2019), who investigated diploid *Coffea* species. The combination of GBS with a multilabeled (MUL) tree, *i.e.* a phylogenetic model wherein the subgenomes of hybrid species are displayed as separate tips as if they would be distinct species (Huber *et al*., 2006), is a promising approach to investigate the evolutionary origin of the allotetraploid *C. arabica*. Analyzing hybrid evolution via a MUL tree has the advantage over a phylogenetic network analysis in that it allows for divergence time analyses to estimate the age of a hybridization event (Estep *et al*., 2014; Marcussen *et al*., 2015; McCann *et al*., 2018).

Here, we aim to infer the evolutionary origin of the self-compatible allotetraploid species *C. arabica* based on GBS genome fingerprinting combined with a MUL tree approach. We included 23 *Coffea* species among which the seven species that are known to be closely related to *C. arabica.* Our research questions were: (i) Which extant *Coffea* species are genetically most closely related to *C. arabica*? (ii) When did the hybridization event at the origin of *C. arabica* occur? (iii) How did self-compatibility evolve in the *Coffea* genus?

## Materials & methods

### Taxon sampling and DNA extraction

Leaf samples were collected from 35 accessions of *Coffea* and one of *Tricalysia*, the latter serving as outgroup (Table 1, Supporting Information Table S1). All accessions were part of the herbarium (BR) and the living collections of Meise Botanic Garden, Belgium. The ingroup encompassed 23 *Coffea* species with at least one representative of each of the main clades in the phylogeny of the genus (Hamon *et al*., 2017; Guyeux *et al*., 2019). All known species that are closely related to *C. arabica* (*i.e. C. brevipes, C. canephora, C. congensis, C. anthonyi, C. eugenioides, C. heterocalyx*, and *C. kivuensis*) were also part of the ingroup (Maurin *et al*., 2007). DNA extractions were carried out using an optimized cetyltrimethylammonium bromide (CTAB) protocol adapted from Doyle & Doyle (1987). DNA quantities were measured with the Quantifluor dsDNA system on a Promega Quantus Fluorometer (Promega, Madison, USA).

**Table 1.** Overview of the species and the number of accessions that were included in this study.

| **Taxon name** | **Number of accessions** |
| --- | --- |
| Outgroup |  |
| *Tricalysia* sp. | 1 |
| Ingroup |  |
| *Coffea anthonyi* Stoff. & F.Anthony | 2 |
| *Coffea arabica* L. | 3 |
| *Coffea brevipes* Hiern | 2 |
| *Coffea canephora* Pierre ex A.Froehner | 2 |
| *Coffea charrieriana* Stoff. & F.Anthony | 2 |
| *Coffea congensis* A.Froehner | 2 |
| *Coffea dubardii* Jumelle | 1 |
| *Coffea eugenioides* S.Moore | 2 |
| *Coffea humilis* A.Chev. | 1 |
| *Coffea kapakata* (A.Chev.) Bridson | 1 |
| *Coffea kivuensis* Lebrun | 2 |
| *Coffea lebruniana* Germ. & Kesler | 1 |
| *Coffea liberica* ex Hiern | 2 |
| *Coffea lulandoensis* Bridson | 2 |
| *Coffea macrocarpa* A.Rich. | 1 |
| *Coffea mannii* (Hook.f.) A.P.Davis | 1 |
| *Coffea pocsii* Bridson | 1 |
| *Coffea pseudozanguebariae* Bridson | 1 |
| *Coffea racemosa* Lour. | 1 |
| *Coffea salvatrix* Swynn. & Philipson | 1 |
| *Coffea sessiliflora* Bridson | 1 |
| *Coffea stenophylla* G.Don | 2 |
| *Coffea heterocalyx* Stoff. | 1 |

### GBS Library preparation

GBS libraries were prepared using a single-enzyme protocol that was slightly adapted from Elshire *et al*. (2011). 100 ng of DNA was digested with *Pst*I (New England Biolabs, Ipswitch, USA). The digested DNA fragments were ligated to a barcode adapter-common adapter system (0.045 pmol) with T4 DNA ligase (New England Biolabs, Ipswitch, USA). Each in-line barcode was between four and nine basepairs (bp) long, differed from all other barcodes by at least three sites, and had no homopolymers longer than 2 bp. Ligation products were purified with 1.6X MagNa magnetic beads (Rohland & Reich, 2012) and eluted in 30 µl TE. Of the purified DNA eluate, 3 µl were used for amplification with *Taq* 2X Master Mix (New England Biolabs, Ipswitch, USA) using a 20-cycles PCR protocol. PCR products were bead-purified with 1.6X MagNa, and their DNA concentrations were quantified with the Quantus Fluorometer. Afterwards, fragment size distributions were assessed using a Qiagen QIAxcel system (Qiagen, Venlo, NL). Equimolar amounts of the GBS libraries were pooled, bead-purified, and 150 bp paired-end sequenced on an Illumina HiSeq-X instrument by Admera Health (South Plainfield, USA). Technical GBS library replicates of 13 samples were made to test reproducibility of genetic distance estimates.

### Data processing

The quality of sequence data was validated with FastQC v0.11 (Andrews, 2010) and reads were demultiplexed using GBSX v1.3 (Herten *et al*., 2015) with one mismatch allowed in barcodes. The maximum length of forward reads was adjusted to 142 bp in order to compensate for variable barcode lengths. The 3’ restriction site remnant and the common adapter sequence of forward reads and the 3’ restriction site remnant, the barcode, and the barcode adapter sequence of reverse reads were removed with Cutadapt v1.9 (Martin, 2011). The 5’ restriction site remnant of forward and reverse reads was trimmed with FASTX-Toolkit v0.0.13 (Gordon & Hannon, 2010). Next, forward and reverse reads with a minimum read length of 60 bp and a minimum overlap of 10 bp were merged using PEAR v0.9.8 (Zhang *et al*., 2014). Merged reads with a mean base quality below 25 or with more than 5% of the nucleotides uncalled were discarded using prinseq-lite v0.20.4 (Schmieder & Edwards, 2011). Reads containing internal restriction sites were discarded using the OBITools package (Boyer *et al*., 2016). The trimmed sequencing data is available in the NCBI sequence read archive (BioProject PRJNA612193).

### Clustering analyses and genetic distance calculation

Preprocessed reads were analyzed with the GIbPSs toolkit (Hapke & Thiele, 2016), a software package that clusters GBS reads into loci without using a reference genome and that allows for variant calling in mixed-ploidy data (Supporting Information Method S1). We defined a locus as a cluster of at least 20 reads of the same length that consisted of one or more alleles (∼haplotypes). Alleles were sequence variants that were supported by at least 5 identical reads in at least one sample. If the number of nucleotide differences between alleles was less than 10% of the allele length, they were assigned to the same locus. To remove possible contamination, an additional BLAST search against a local reference database was performed for all alleles in the dataset. This database consisted of RefSeq genomes of viruses, prokaryotes, and fungi (O’Leary *et al*., 2016), the reference genome sequence of *C. canephora* (Denoeud *et al*., 2014), and the reference chloroplast genome sequence of *C. arabica* (Genbank accession number NC_008535.1). At the time of our analyses, the *C. canephora* reference genome was the only published high quality reference genome sequence of a *Coffea* species. As the *C. canephora* chloroplast reference genome sequence was found to differ substantially from the chloroplast genome sequence of many other *Coffea* species (Guyeux *et al*., 2019), the use of the *C. arabica* chloroplast reference genome sequence resulted in the identification of additional chloroplast alleles in our dataset. The expect value cutoff of the BLAST search was set to 0.0001 and the single best search result was used to deduce the putative origin of each allele. Loci with no allele matching the *Coffea* reference genome sequences were removed. Next, loci with data in less than 5% of the samples were discarded to reduce the amount of missing data. Jaccard genetic similarities (J) between pairs of samples were calculated as the number of common alleles divided by the total number of alleles (Jaccard, 1912). Only loci without missing data in both samples were taken into account for these calculations. Genetic distances were subsequently calculated as 1 – J and visualized in a genetic distance matrix using a custom python script.

### Phylogenetic and divergence time analyses

One representative sample of each species was selected for MUL phylogenetic tree reconstruction to reduce computation time of the analysis. As genetic distances between conspecific samples were acceptably low compared to interspecific genetic distances, reducing the number of samples per species did not influence the results. Loci of *C. arabica* samples were split into two distinct sets, which corresponded to the subgenomes of the *C. arabica* genome, following a two-step approach. First, the alleles of the putative progenitor species of *C. arabica* were partitioned into two groups: one group with all alleles of *C. brevipes, C. canephora*, and *C. congensis* and another with all alleles of *C. anthonyi, C. eugenioides, C. kivuensis*, and *C. heterocalyx*. Next, each *C. arabica* allele was compared to all alleles of the same locus in both groups of progenitor species. If a *C. arabica* allele was more similar to an allele in one group of progenitors than to any of the alleles in the other group, the allele was assigned to the corresponding “group-specific” subgenome of *C. arabica*. The entire locus was discarded if allelic data was absent in one or both progenitor groups or if not all alleles of that locus could be assigned to one of the two progenitor groups. Afterwards, locus data was converted into consensus alignments using custom python scripts. The scripts used to process the GIbPSs output files are available on GitLab (https://gitlab.com/ybawin/origin_coffea_arabica).

Next, the most optimal substitution model was determined for each locus alignment based on the Akaike Information Criterion corrected for small sample size (AICc) using jModelTest v2.1.10 (Darriba *et al*., 2012). A Maximum Likelihood multilabeled (MUL) phylogenetic tree was reconstructed for each locus alignment with RAxML v8 (Stamatakis, 2014). Up to 1000 thousand bootstrap replicates were created for each alignment, but bootstrapping was halted when support values stabilized earlier, which was tested using the extended majority-rule consensus tree criterion (Pattengale *et al*., 2010). A 75% majority-rule consensus tree was reconstructed for each locus and the information in all locus trees was summarized in one consensus tree using ASTRAL-III (Zhang *et al*., 2018). A local posterior probability (localPP) threshold of 0.95 was used to accept nodes.

In addition, a Bayesian MUL phylogenetic tree was reconstructed for each locus alignment with MrBayes v3.2.6. (Ronquist *et al*., 2012). The number of generations per run was set to five million. A relative burn-in of ten percent was applied and three replicate runs were performed for every alignment. Convergence within each run was assessed based on the effective sampling size (ESS) (> 200) and the potential scale reduction factor (between 0.99 and 1.01). Convergence between runs was evaluated using the average standard deviation of split frequencies (< 0.01). A consensus tree was reconstructed based on all locus trees using ASTRAL-III. Nodes with a localPP lower than 0.95 were removed.

A Bayesian Markov Chain Monte Carlo (MCMC) divergence time analysis was done with BEAST v1.10 (Suchard *et al*., 2018) and parameters for this analysis were set in BEAUTi v1.10 (Suchard *et al*., 2018). GBS data of separate loci were concatenated to reduce computational complexity. The most parameter-rich substitution model (GTR+G+I) was chosen for the entire dataset, which should compensate for deviations from this model for separate loci and provide accurate results (Abadi *et al*., 2019). The age of the most recent common ancestor of the *C. mannii* - *C. lebruniana* clade and all other *Coffea* species was re-estimated using the age estimate of Tosh *et al*. (2013) as secondary calibration point and a normal prior (mean = 10.77 Ma, SD = 1.0 Ma). Evolution was modeled as a Yule process and rates varied across lineages according to an uncorrelated relaxed lognormal molecular clock (Drummond *et al*., 2006). This clock model was selected based on marginal likelihood estimations using the generalized stepping-stone sampling method (Baele *et al*., 2016) with five hundred stepping stones and a chain length of one million generations. Five hundred million generations were run to complete the analysis with trees sampled every five thousand generations. Three replicate runs were performed and chain convergence, run convergence, and ESS parameter estimation (> 200) were evaluated with Tracer v1.7.1 (Suchard *et al*., 2018). The results of the three runs were combined with LogCombiner v1.10 (Suchard *et al*., 2018) and a maximum clade credibility tree with a posterior probability limit of 0.9 was reconstructed using TreeAnnotator v1.10.1 (Suchard *et al*., 2018).

The evolution of self-compatibility in *Coffea* was inferred using the BEAST maximum clade credibility tree and a Maximum Likelihood reconstruction method implemented in Mesquite v2.75 (Maddison & Maddison, 2006, 2011). The ability of species to self-pollinate was coded as a binary trait (0 = self-incompatible (SI), 1 = self-compatible (SC)) and likelihoods were calculated using a Markov *k*-state one-parameter model (Mk1), assuming a single transition rate between SI and SC. Character states were assigned to nodes based on a likelihood ratio test with a likelihood decision limit of two. If the difference in log-likelihood of SI and SC was two or more, the state with the highest likelihood was accepted as the most likely state. Nodes with a log-likelihood difference lower than two were considered to be ambiguous.

## Results

### GBS summary data and ancestry of Coffea arabica

In total, 23 676 loci (including 47 chloroplast loci) of a size between 60 and 273 bp and with data for at least 5 percent of the samples were retrieved. Out of a total of 3 901 029 nucleotide sites sequenced, 237 619 sites (6.09 %) were variable. The number of loci without missing data in each pair of samples was sufficiently high to obtain stable genetic distance estimates (Fig. 1a, Supporting Information Fig. S1, Supporting Information Fig. S2). However, the number of chloroplast loci was too low to infer distance estimates solely based on this set of loci (data not shown). Genetic distance values were highly reproducible, as genetic distances between technical replicates (0.02 – 0.04) were much lower than the mean genetic distance between different accessions (0.88) (Fig. 1b, Supporting Information Fig. S3). Considering the two species groups containing all species closely related to *C. arabica*, genetic distances between *C. brevipes, C. canephora*, and *C. congensis* on the one hand and *C. anthonyi, C. heterocalyx, C. eugenioides*, and *C. kivuensis* on the other were moderately high (0.87 – 0.89). Within these groups, *C. eugenioides* (0.66 – 0.68) and *C. canephora* (0.63 – 0.65) displayed the lowest genetic distances to the *C. arabica* accessions (Fig. 1b, Supporting Information Fig. S3). These distances were substantially lower than the genetic distance between *C. arabica* and the second-most genetically similar species in each group (*C. kivuensis*: 0.73 – 0.76; *C. congensis*: 0.78-0.79), showing that among the species included in this analysis, *C. eugenioides* and *C. canephora* are genetically most closely related to *C. arabica*.

**Fig. 1.**
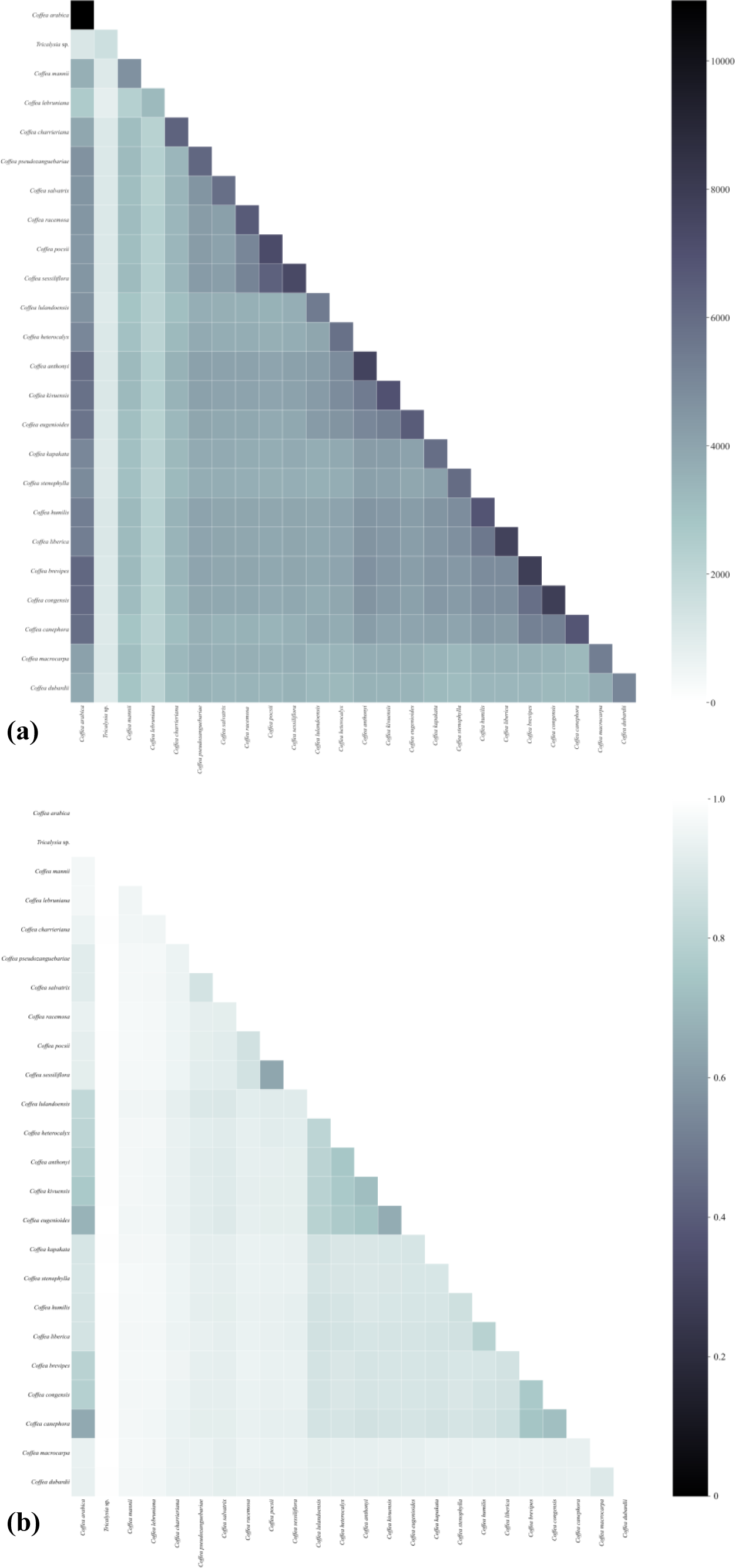
Heat map of the number of common loci **(a)** and the Jaccard genetic distance estimates **(b)** between the 23 *Coffea* accessions and one *Tricalysia* accession that were included in the molecular dating analysis. Annotated heat maps of the complete data set are available as supporting information (supporting information Fig. S2 and S3). The number of common loci is indicated in false color ranging from white (no common loci) to black (11 thousand common loci). The Jaccard genetic distance estimates are shown in false color ranging from black (identical) to white (completely different). *Coffea eugenioides* and *C. canephora* were genetically most similar to *C. arabica*, confirming that they are the putative progenitors of this species.

### Phylogenetic reconstruction and divergence time analysis

The topology of the phylogenetic trees reconstructed with Maximum Likelihood (Supporting Information Fig. S4) and Bayesian inference were identical (Supporting Information Fig. S5). Within these trees, *C. arabica* subgenome A was sister to *C. eugenioides* (localPP: 1), whereas *C. arabica* subgenome B was sister to *C. canephora* (localPP: 1). In the clade of *C. arabica* subgenome A, the West-African species *C. anthonyi* and *C. heterocalyx* branched off first followed by *C. kivuensis*, which was positioned close to *C. eugenioides.* In the clade containing *C. arabica* subgenome B, *C. brevipes* was the most early diverged species, while *C. congensis* was a sister species to *C. canephora.* The stem age of the *C. arabica* subgenome A was estimated around 934 thousand years, while the stem age of the *C. arabica* subgenome B was estimated around 720 thousand years (Fig. 2). The highest posterior density interval of both estimates, which is the credible interval containing 95% of the values sampled by the MCMC chain, overlapped between 543 thousand years and 1.08 million years. The age estimates of *C. arabica* were much younger than the estimates of other *Coffea* species in our dataset.

**Fig 2.**
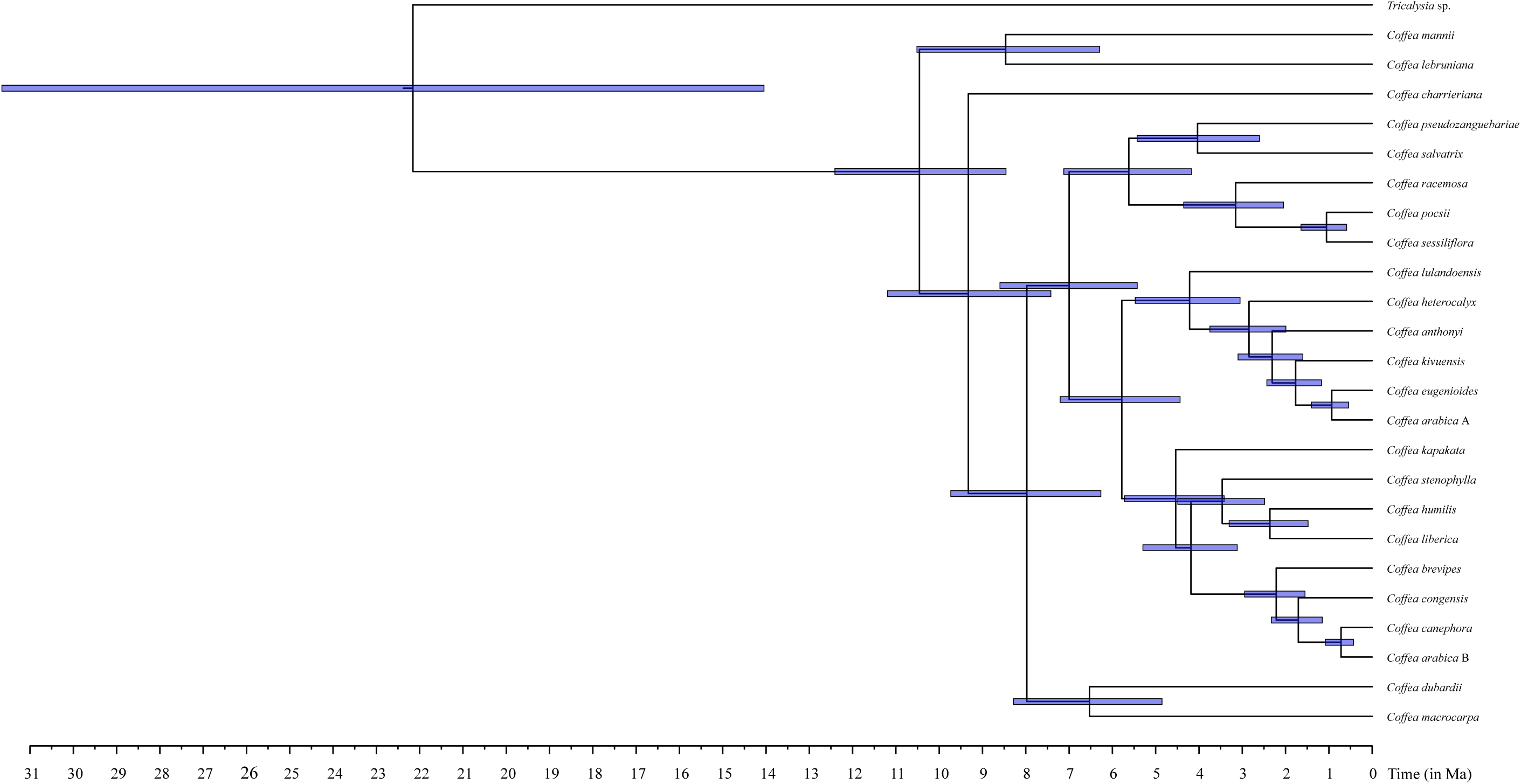
BEAST maximum clade credibility tree of the genus *Coffea* inferred from a combined dataset of variable GBS loci comprising 551 852 nucleotide sites. A *Tricalysia* species was used as outgroup. The node corresponding to the secondary calibration point is indicated by an asterisk (*). Blue node bars display highest posterior density (HPD) intervals and a time scale axis (in Ma) is depicted below the tree. HPD intervals of the *C. arabica* subgenomes overlapped between 543 thousand years and 1.08 million years ago, situating the origin of *C. arabica* within this time interval.

The ancestral state reconstruction of self-compatibility in *Coffea* showed that most ancestors of extant *Coffea* species were most likely self-incompatible (Supporting Information Fig. S6). The ancestral state of all nodes in the clade comprising *C. arabica* subgenome A and the two other known self-compatible *Coffea* species (*i.e. C. heterocalyx* and *C. anthonyi*) remained ambiguous, as SI and SC were both assigned to the recent common ancestors of these species with similar log likelihood values.

## Discussion

The current study provides a clear hypothesis regarding the evolutionary origin of *C. arabica*. GBS data proved to be more informative than the molecular data used in previous studies (Lashermes *et al*., 1997; Cros *et al*., 1998; Raina *et al*., 1998; Lashermes *et al*., 1999; Maurin *et al*., 2007; Tesfaye *et al*., 2007; Hamon *et al*., 2009), also because a substantial amount of informative sites seems to be required to get reliable genetic distance estimates for *Coffea* species (Fig. 1). Based on the similarity in plastid DNA markers, previous research suggested that *C. eugenioides* or a close relative of this species was the ovule donor in the *C. arabica* hybridization event (Maurin *et al*., 2007; Tesfaye *et al*., 2007; Guyeux *et al*., 2019). In this study, we confirmed that *C. eugenioides* is genetically more similar to *C. arabica* than *C. anthonyi, C. heterocalyx*, and *C. kivuensis. Coffea kivuensis* was positioned as a sister species to *C. eugenioides* and *C. arabica* subgenome A (Fig. 2, Supporting Information Fig. S4, Fig. S5), corroborating the high morphological and ecological similarity between *C. kivuensis* and *C. arabica*. Chevalier (1947) classified *C. kivuensis* as a variety of *C. eugenioides*, which is in accordance with the low genetic distances between *C. kivuensis* and *C. eugenioides* found in this study. However, we believe that the classification of *C. kivuensis* as a separate species is justified, because genetic distance values between these species were substantially higher than the intraspecific genetic distances estimated in this study (Fig. 1, Supporting Information Fig. S3). The West-Central African species *C. anthonyi* and *C. heterocalyx* were more distantly related to *C. arabica*, reflecting their geographical distance to this species.

Using our GBS and MUL tree approach, we confirmed that *C. canephora* was the putative pollen donor in the hybridization event prior to the emergence of *C. arabica. C. congensis* and *C. brevipes* were clearly more distantly related to *C. arabica* subgenome B (Fig. 2, Supporting Information Fig. S4, Fig. S5). *Coffea canephora* currently has one of the widest natural distribution ranges in the *Coffea* genus, reaching from Guinea to Tanzania, but it does not naturally co-occur with *C. arabica* (Supporting Information Fig. S7, Davis *et al*., 2006). Using single nucleotide polymorphisms in *C. canephora* individuals that were sampled across its entire natural range, *C. arabica* was found to be genetically most similar to *C. canephora* accessions in northern Uganda (Merot-L’anthoene *et al*., 2019; Scalabrin *et al*., 2020).

Although the natural ranges of *C. canephora* and *C. eugenioides* overlap in East-Central Africa (Supporting Information Fig. S7), natural hybrids between these species in this area are not known so far. The absence of known recent hybrids between *C. canephora* and *C. eugenioides* can be explained by three factors. First, although both species can be found in the same area, their habitat preference differs substantially. *Coffea eugenioides* is especially found near forest edges, while *C. canephora* is mainly restricted to the forest interior (Noirot *et al*., 2016). Second, the flowering time of both species does not coincide (Noirot *et al*., 2016). The flowering time of *Coffea* species is highly species-specific and genetically controlled, hampering interspecific gene flow via pollination (Gomez *et al*., 2016). Third, the success rate of induced cross-pollination between *C. canephora* and *C. eugenioides* is very low, suggesting the presence of additional reproductive barriers (Noirot *et al*., 2016). However, changes in environmental conditions may have broken (some of the) reproductive barriers between *C. canephora* and *C. eugenioides* in the past, enabling a successful interspecific hybridization between these species at the origin of *C. arabica.* In support of this hypothesis, Gomez *et al*. (2016) reported that the flowering time of *C. arabica, C. canephora*, and *C. liberica* became more synchronized in New Caledonia, in response to changes in precipitation regime, resulting in the emergence of spontaneous hybrids. Similar events were also observed in living collections, where different *Coffea* species that were *ex situ* conserved in a common environment more easily hybridized (Noirot *et al*., 2016).

We estimated the time of the *C. arabica* hybridization event between 1.08 million and 543 thousand years ago. The *C. arabica* subgenomes were the youngest taxa within the phylogenetic tree, meaning that the diversity in extant *Coffea* species was generally established before *C. arabica* emerged. The age interval found in this study overlaps with the maximum age estimate of Yu *et al*. (2011) but contained much older estimates than the estimates provided by Cenci *et al*. (2012) and Scalabrin *et al.* (2020). Interestingly, age estimates based on the diversity *within C. arabica* (Cenci *et al*., 2012; Scalabrin *et al*., 2020) situated the origin of *C. arabica* much more recent than estimates based on the diversity *between Coffea* species (Yu *et al*., 2011; this study), which might suggest that *C. arabica* underwent a large genetic bottleneck (Scalabrin *et al*., 2020). Moreover, the median values of the stem age estimates of both subgenomes (*i.e.* 965 and 720 thousand years) were higher than the estimate of Yu *et al.* (2011), plausibly situating the *C. arabica* hybridization event further back in time. Changing environmental conditions during this period might have played an important role in *C. arabica* speciation. Pollen records of marine and lake sediment cores in the Congo basin and East Africa indicate that the Afromontane forest regularly expanded to lower altitudes during the glacial periods between 1.05 million and 600 thousand years ago (Dupont *et al*., 2001; Owen *et al*., 2018). These forest expansions might have enlarged the contact zone between *C. canephora* and *C. eugenioides* and the coinciding altered environmental conditions may have weakened interspecific reproductive barriers between these species. Moreover, the aridification of East-Africa over the past 575 thousand years may have changed the natural ranges of *C. canephora, C. eugenioides*, and *C. arabica* to their current distribution area (Owen *et al*., 2018). The emergence of hybrid species is often linked to climate-induced range shifts of progenitor species (Kadereit, 2015; Arnold, 2016; Wagner *et al*., 2019). Likewise, the origin and subsequent emergence of *C. arabica* might have been influenced by climate fluctuations in East-Africa during the last one million years.

The reconstruction of the evolution of self-compatibility in *Coffea* showed that the character state regarding self-compatibility of each node in the clade of *C. arabica* subgenome A and other self-compatible *Coffea* species (*C. heterocalyx* and *C. anthonyi*) could not unambiguously be inferred (Supporting Information Fig. S6). However, the fact that *C. arabica* is closest related to two self-incompatible species may suggest that the ovule donor of *C. arabica* was self-incompatible as well. Consequently, self-compatibility in *Coffea* most likely evolved first in the most recent common ancestor of *C. heterocalyx* and *C. anthonyi*, followed by a reversal to self-incompatibility in the most recent common ancestor *C. kivuensis* and *C. eugenioides*, and the independent development of self-compatibility in *C. arabica*. The breakdown of self-incompatibility in allopolyploids with self-incompatible progenitor species is believed to be a survival strategy to assure reproduction when the number of available mating partners is limited (Osabe *et al*., 2012). The presence of self-compatible species at the basis of the clade containing *C. arabica* subgenome A may suggest that the ancestor of *C. arabica* possessed a certain aptitude for the change to self-compatibility that may have facilitated its survival after its emergence.

Age estimates of allopolyploids may deviate from their actual age because the genotypes of progenitor species included in the dating analyses were divergent from the actual progenitor genotypes (Doyle & Egan, 2010). Our *C. canephora* specimens were sampled in D.R. Congo from populations closely related to the Ugandan populations (Supporting Information Table S1), which were found to be genetically most similar to *C. arabica* (Merot-L’anthoene *et al*., 2019; Scalabrin *et al*., 2020). Although we do not know which *C. eugenioides* populations are genetically closest to *C. arabica*, age estimates of *C. arabica* are probably less affected by the origin of the *C. eugenioides* genotype as genetic diversity in this species was found to be very low compared to the diversity in *C. canephora* (Merot-L’anthoene *et al*., 2019).

Overall, we have clearly confirmed *C. canephora* and *C. eugenioides* as the closest known relatives of *C. arabica.* The hybridization event at the origin of *C. arabica* was estimated between 1.08 million and 543 thousand years ago and was linked to changing environmental conditions in East-Africa during glacial-interglacial cycles in the last one million years. We inferred that self-compatible species in *Coffea* were a paraphyletic group and that self-compatibility most likely evolved twice in *Coffea.* Our research clarified the evolutionary relationships between the direct wild relatives of cultivated Arabica coffee, providing a strong instrument for the selection of wild plant species in coffee breeding programs. The closest relatives of a crop often contain more favorable characteristics for breeding than distantly related species, showing the importance of phylogenetic studies on crop wild relatives for crop improvement (Preece *et al*., 2015, 2018; Martín-Robles *et al*., 2019).

## Acknowledgements

We thank Cephas Masumbuko Ndabaga, Eberhard Fischer, Samuel Vanden Abeele, and Filip Vandelook for collecting plant material, and Sabine Van Glabeke and Zyon Vansteelandt for their technical support. This research was funded by Research Foundation - Flanders (FWO) (project N° G056517N).

## Author Contribution

IRR, OH, and SJB designed the research. JCIMM and PS provided the leaf material. YB, TR, and AS planned and executed the lab work. YB, TR, AH, and SJB processed and analyzed the sequence data. YB wrote the manuscript, which was revised and commented by all authors.

