## Supplementary materials for "Phylogenomic analysis clarifies the evolutionary origin of *Coffea arabica* L"

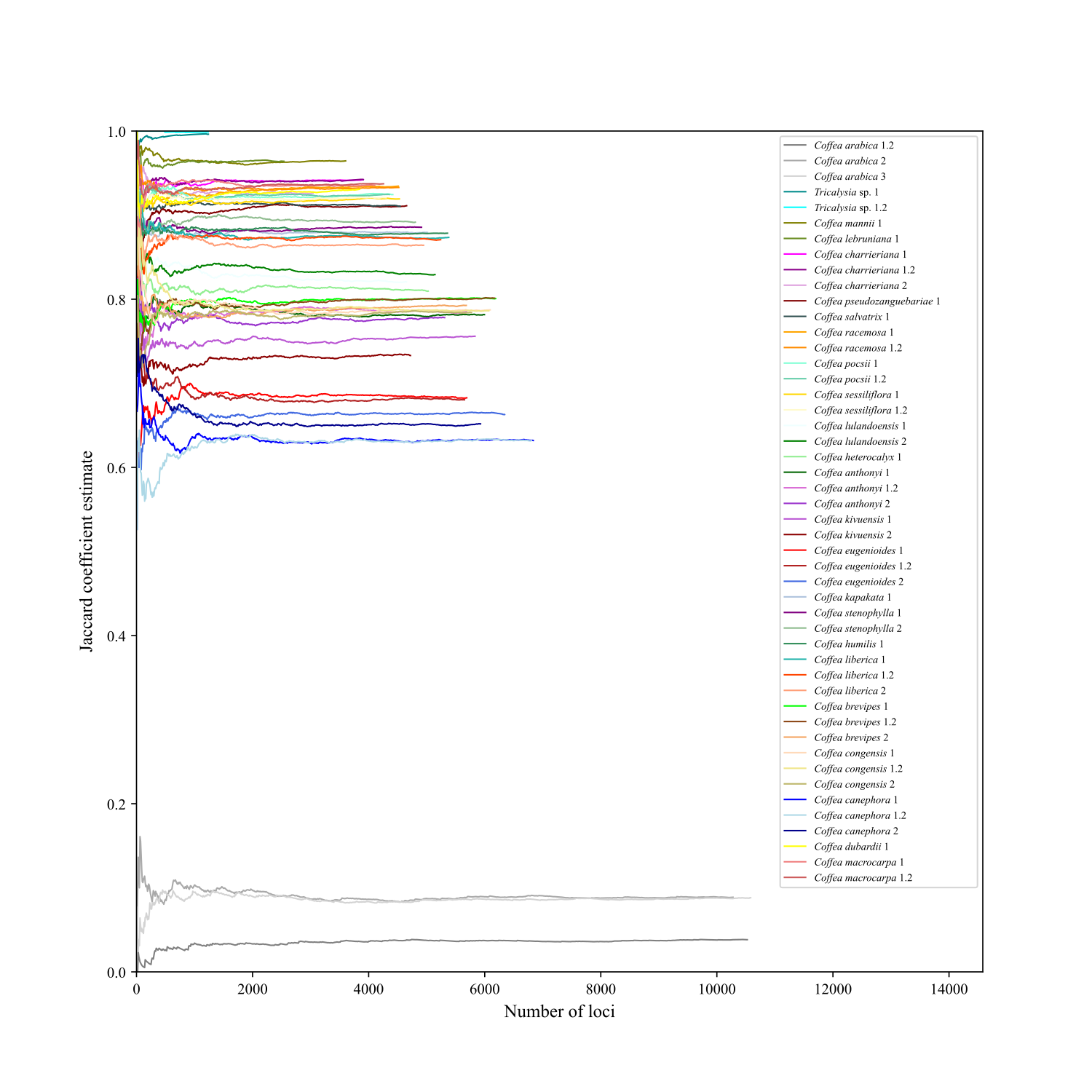

**Fig. S1** Line plot showing genetic distance estimates between sample ‘*Coffea arabica* 1’ and all other samples calculated after every ten loci. The x-axis displays the number of loci, the y-axis genetic distance estimates. The number of common loci in each comparison is sufficiently high to obtain stable genetic distance estimates.

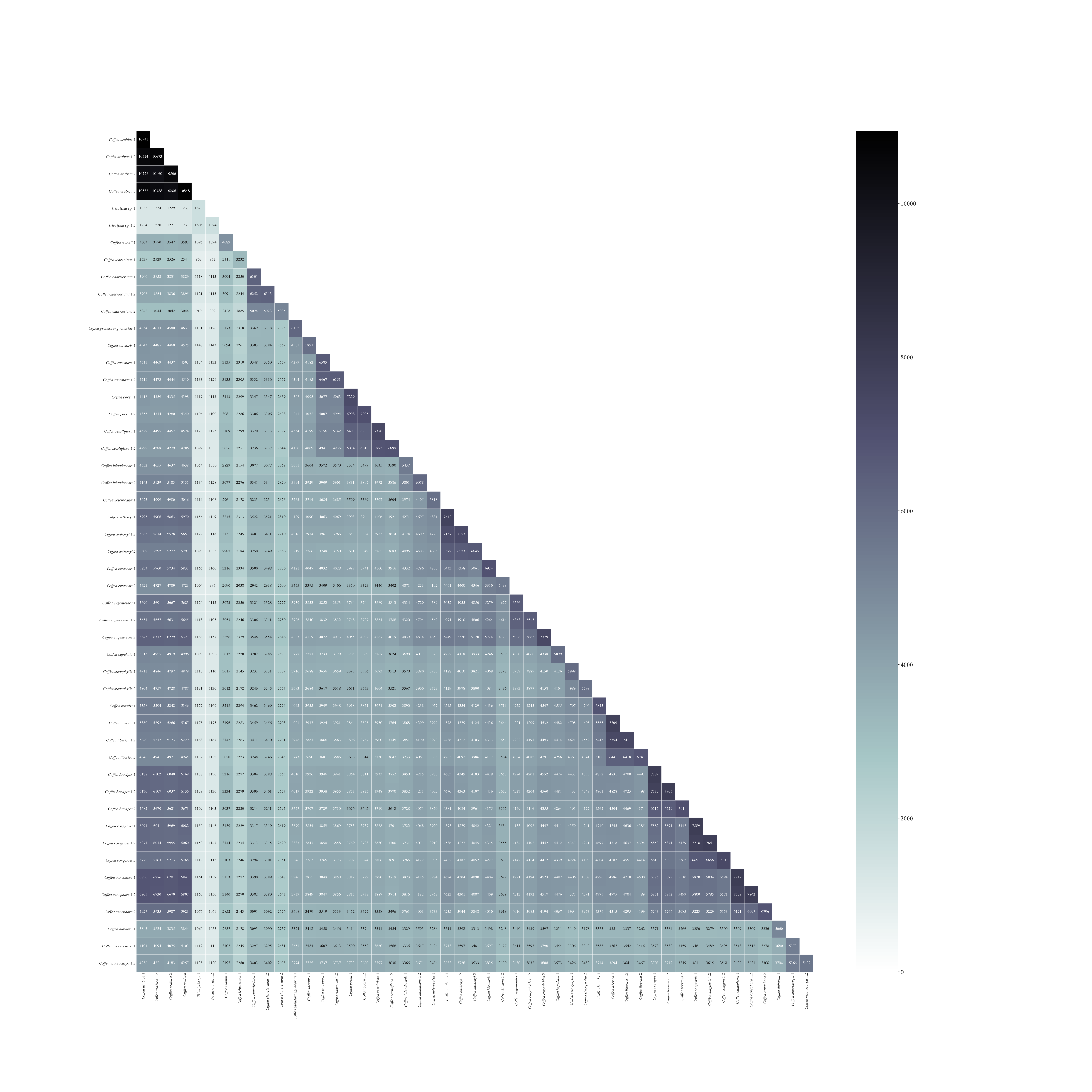

**Fig. S2** Annotated heat map of the number of common loci between 35 accessions of *Coffea* and one of *Tricalysia*. Technical replicates have the same name as the original sample followed by the suffix ‘.2’. Numbers are indicated in false color ranging from white (no common loci) to black (11 thousand common loci).

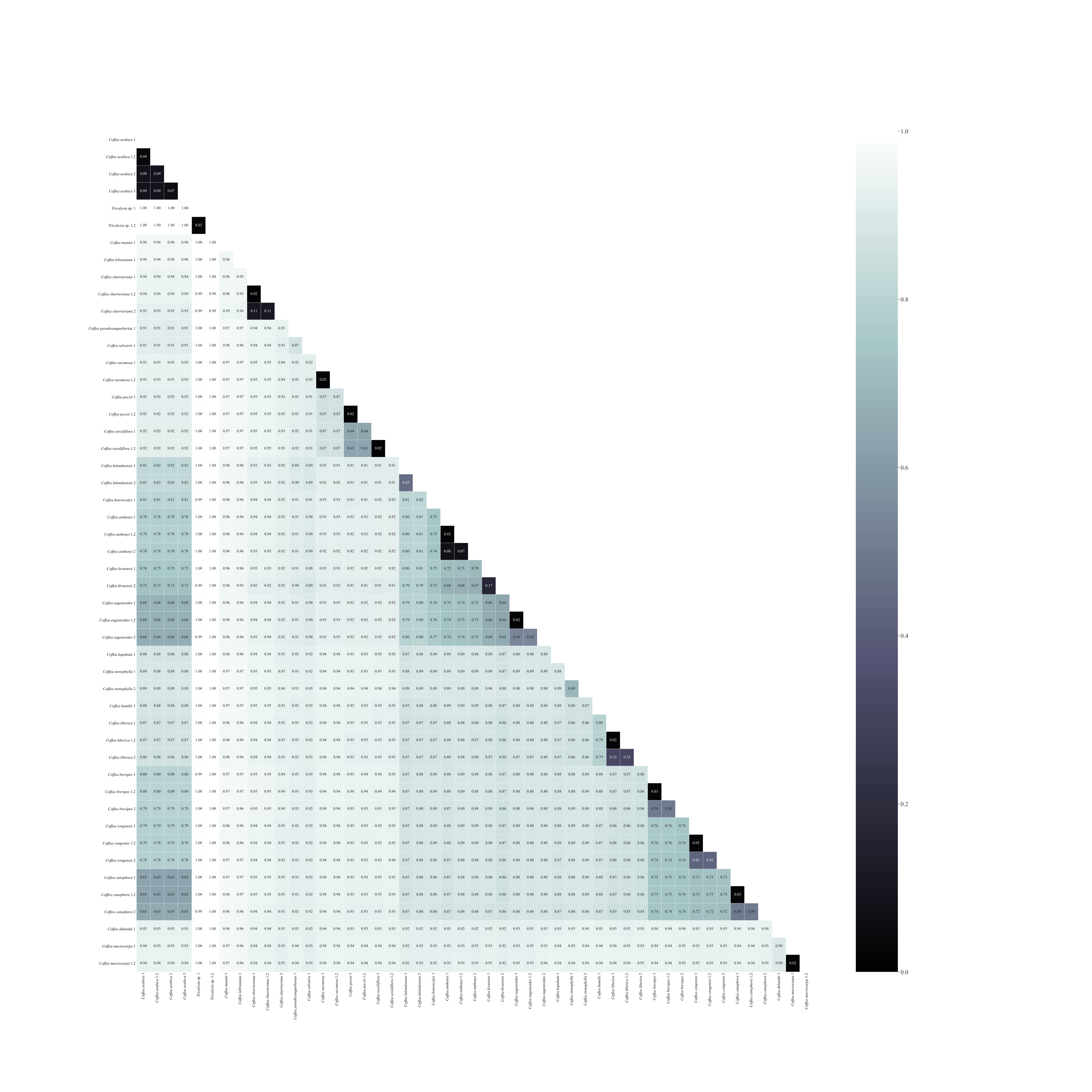

**Fig. S3** Annotated heat map of pairwise Jaccard genetic distances between 35 accessions of *Coffea* and one of *Tricalysia*. Technical replicates have the same name as the original sample followed by the suffix ‘.2’. Genetic distances are indicated in false color ranging from black (identical) to white (completely different).

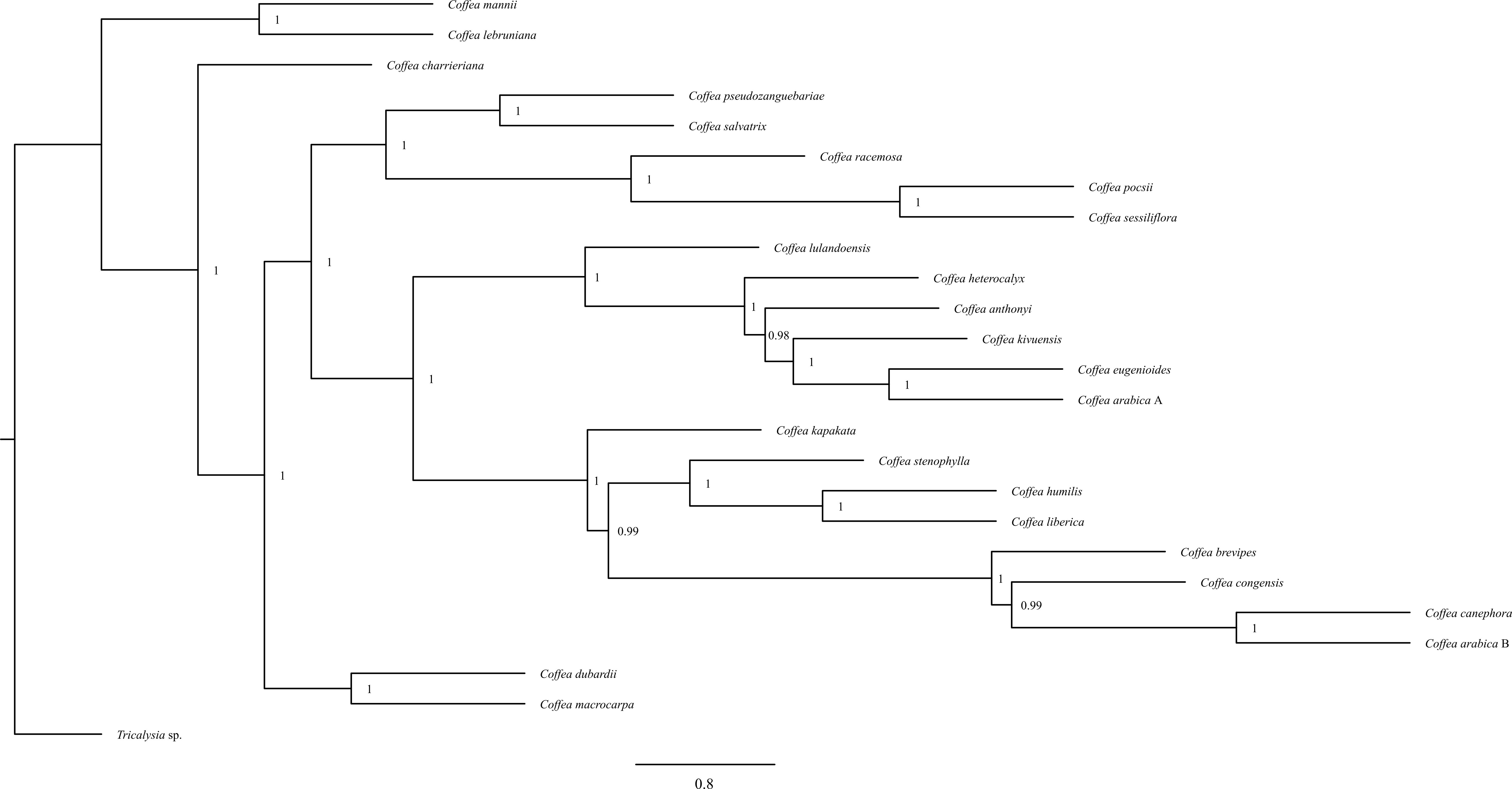

**Fig. S4** ASTRAL consensus tree of 3290 Maximum Likelihood multilabeled locus trees of 24 *Coffea* species and one *Tricalysia* species. Maximum Likelihood trees were reconstructed using 1000 bootstrap replicates and a 75% majority-rule consensus criterion. Node labels indicate local posterior probability values.

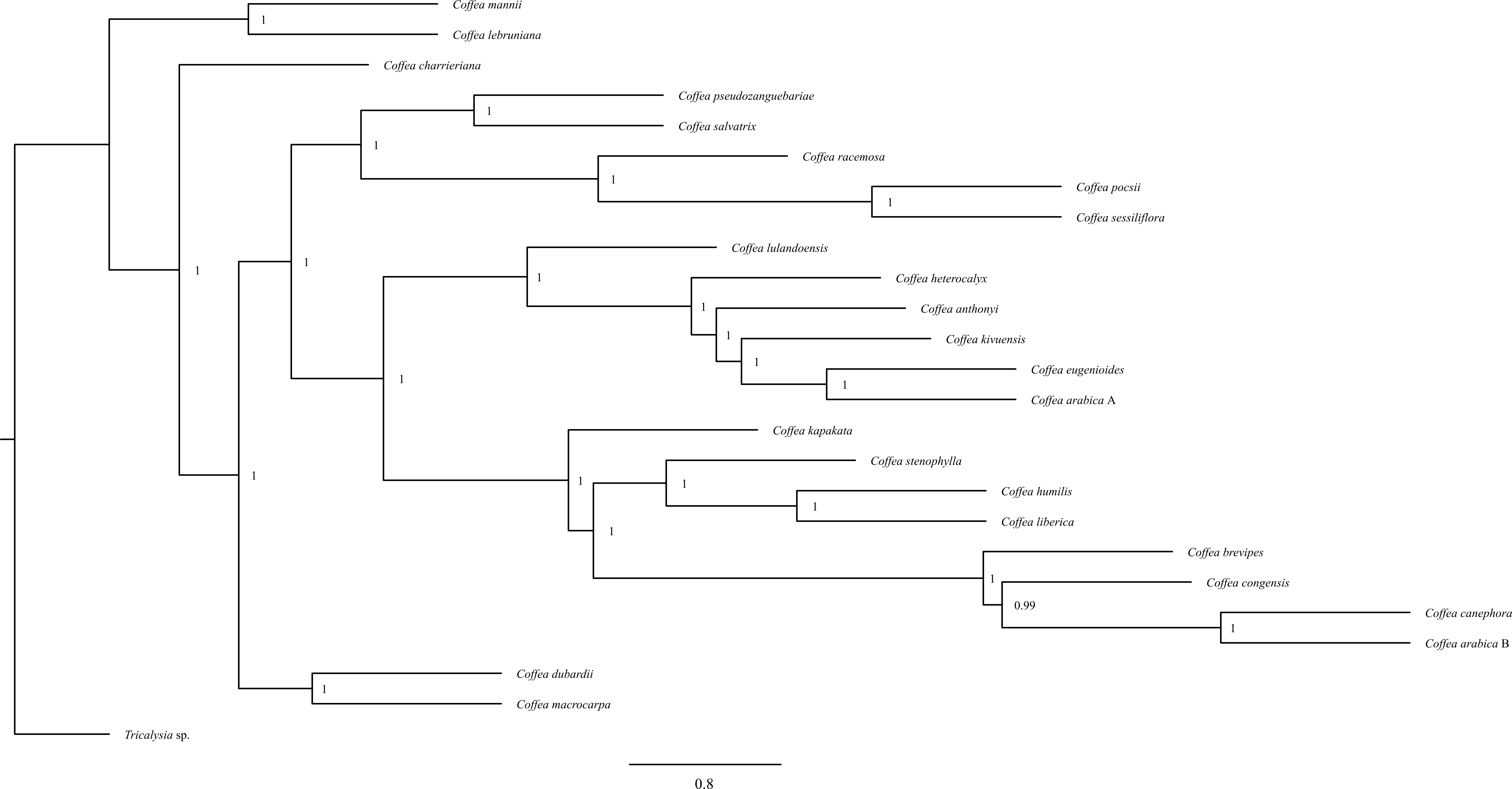

**Fig. S5** ASTRAL consensus tree of 3290 Bayesian inference multilabeled locus trees of 24 *Coffea* species and one *Tricalysia* species. Bayesian inference consensus trees were reconstructed based on 3 replicate runs of 5 million generations. Node labels indicate local posterior probability values.

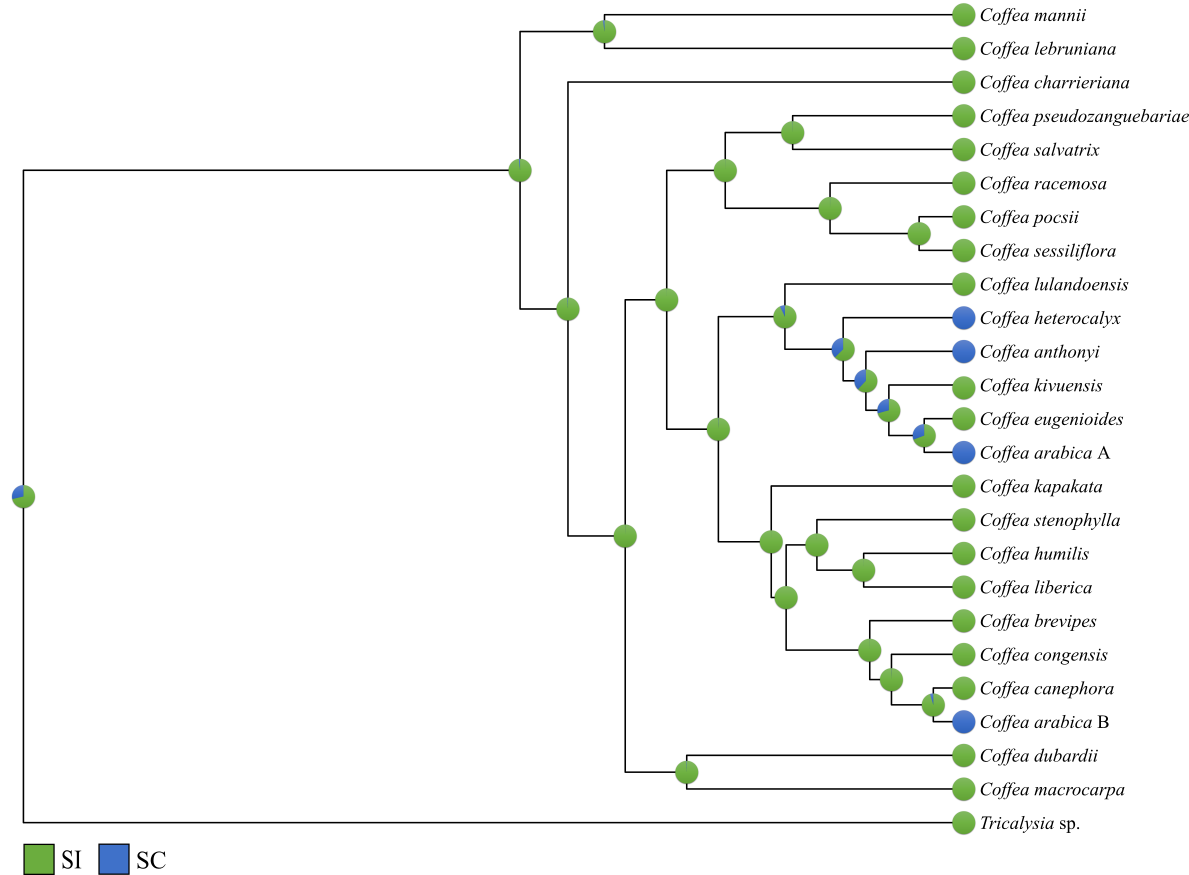

**Fig. S6** Ancestral state reconstruction of self-compatibility in the *Coffea* genus*.* Character states were coded as SI (self-incompatible, green) or SC (self-compatible, blue). Pie charts display the proportional likelihoods of each state, which is the proportion of the raw likelihood value of the state to the sum of the raw likelihood values of both states.

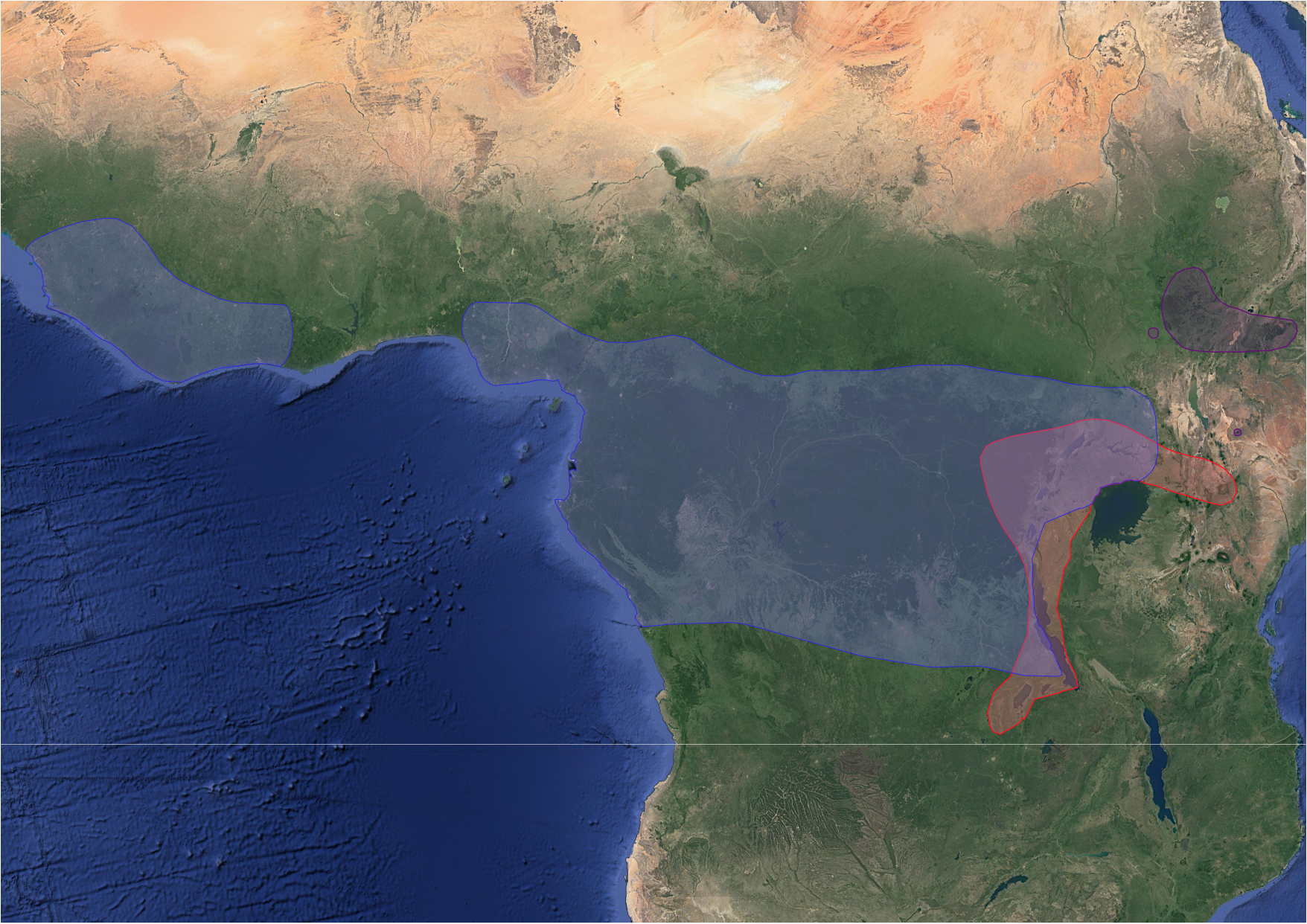

**Fig. S7** Map showing the geographic distribution of *C. arabica* (purple), *C. canephora* (blue), and *C. eugenioides* (red) adapted from Bridson (1982), Bridson & Verdcourt, (1988), Stoffelen (1998), and Davis *et al.* (2012). Map data ©2020 Google.

**Table S1** List of plant accessions used for this study. All accessions originated from Meise Botanic Garden. The taxon name, accessions number/universally unique identifier, country of origin, and accession type (LC: living collection, H: herbarium, SC: silica-dried leaves collection) are listed for each accession. Accessions that were included in the molecular dating analysis are marked in the last column.

| **Taxon name** | **Accession number/universal unique identifier** | **Country** | **Type** | **Molecular dating** |
| --- | --- | --- | --- | --- |
| Outgroup | | | | |
| *Tricalysia* sp. | 20121180-82 | Tanzania | LC | Yes |
| Ingroup | | | | |
| *Coffea anthonyi* 1 | 20070347-77 | Cameroon | LC | Yes |
| *Coffea anthonyi* 2 | 20110333-02 | Cameroon | LC | No |
| *Coffea arabica* 1 | 19690431 | Unknown | LC | Yes |
| *Coffea arabica* 2 | 20110257-23 | Ethiopia | LC | No |
| *Coffea arabica* 3 | 20110259-25 | Ethiopia | LC | No |
| *Coffea brevipes* 1 | 20110326-92 | Cameroon | LC | Yes |
| *Coffea brevipes* 2 | 20110327-93 | Cameroon | LC | No |
| *Coffea canephora* 1 | / | D.R. Congo | SC | Yes |
| *Coffea canephora* 2 | 19391725 | D.R. Congo | LC | No |
| *Coffea charrieriana* 1 | 20070349-79 | Cameroon | LC | Yes |
| *Coffea charrieriana* 2 | BR0000008752734 | Cameroon | H | No |
| *Coffea congensis* 1 | 20110264-30 | Central African Republic | LC | Yes |
| *Coffea congensis* 2 | 20110265-31 | Central African Republic | LC | No |
| *Coffea dubardii* 1 | BR0000009171756 | Madagascar | H | Yes |
| *Coffea eugenioides* 1 | Eberhard, s.n. | Rwanda | SC | Yes |
| *Coffea eugenioides* 2 | BR0000009256408 | Tanzania | H | No |
| *Coffea humilis* 1 | 20110310-76 | Ivory Coast | LC | Yes |
| *Coffea kapakata* 1 | 20110282-48 | Unknown | LC | Yes |
| *Coffea kivuensis* 1 | Masumbuko, 749 | D.R. Congo | SC | Yes |
| *Coffea kivuensis* 2 | BR0000019452418 | D.R. Congo | H | No |
| *Coffea lebruniana* 1 | BR0000019881898 | Gabon | H | Yes |
| *Coffea liberica* 1 | 20110299-65 | Ivory Coast | LC | Yes |
| *Coffea liberica* 2 | 20110298-64 | Ivory Coast | LC | No |
| *Coffea lulandoensis* 1 | BR0000006435417 | Tanzania | H | Yes |
| *Coffea lulandoensis* 2 | BR0000008829627 | Tanzania | H | No |
| *Coffea macrocarpa* 1 | 20110281-47 | Mauritius | LC | Yes |
| *Coffea mannii* 1 | 19770367 | Ghana | LC | Yes |
| *Coffea pocsii* 1 | 20110336-05 | Tanzania | LC | Yes |
| *Coffea pseudozanguebariae* 1 | 20110314-80 | Kenya | LC | Yes |
| *Coffea racemosa* 1 | 20110320-89 | Unknown | LC | Yes |
| *Coffea salvatrix* 1 | 20110331-00 | Tanzania | LC | Yes |
| *Coffea sessiliflora* 1 | 20110359-28 | Kenya | LC | Yes |
| *Coffea stenophylla* 1 | 20131389-09 | Unknown | LC | Yes |
| *Coffea stenophylla* 2 | 20110309-75 | Ivory Coast | LC | No |
| *Coffea heterocalyx* 1 | BR0000006433673 | Cameroon | H | Yes |

**Methods S1** Data analysis workflow with GIbPSs.

Preprocessed reads were analyzed with the GIbPSs toolkit (Hapke & Thiele, 2016, doi: 10.1111/1755-0998.12510), a software package that clusters GBS reads into loci without using a reference genome and allows for variant calling in mixed-ploidy data. Reads were first clustered into alleles with the program Indloc. Reads with less than five copies in the sample (*e.g.* due to read errors) were added to one of the defined alleles if the number of differences was below the predefined maximum allowed number given the length of the read. Afterwards, Single Nucleotide Polymorphisms (SNPs) were called using the frequency threshold method with a minimum frequency of 20%. Alleles with a frequency below 25% after SNP filtering or with sites that only contain Ns were discarded. Next, loci of different samples were clustered into a common locus using the program Poploc if their similarity was higher than 10 percent. After integrating the Indloc and Poploc data with Indpoploc, loci were filtered using Data_selector. Only loci with a length between 60 and 300 hundred bp, without indel mutations, or with no excessive read depth were retained. Loci with aberrant read depths were characterized by a high median scaled depth (the median of the read depths of the considered locus rescaled in regard to the minimal and maximal depth of a locus in all samples) or a low median depth percentage (the median of the percentage of loci with a higher read depth than the considered locus). All loci with a median scaled depth above 0.25 or a median depth percentage below 0.5 were discarded. If alleles of individual loci were assigned to different loci by poploc (*i.e.* split loci), the corresponding individual loci were removed as well. The resulting set of loci was converted into the standard GIbPSs output files that were used for all downstream analyses.
